# Oscillatory mechanisms of intrinsic brain networks

**DOI:** 10.1101/2022.07.07.499235

**Authors:** Youjing Luo, Xianghong Meng, Guangyu Zhou, Jiali Zhou, Guangzhi Deng, Man Li, Jie Xu, Yiman Li, Yue-jia Luo, Hui Ai, Christina Maria Zelano, Fuyong Chen, Pengfei Xu

## Abstract

Neuroimaging studies of hemodynamic fluctuations have shown specific network-based organization of the brain at rest, yet the neurophysiological underpinning of these networks in human brain remain unclear. Here, we recorded resting-state activities of neuronal populations in the key regions of default mode network (DMN, posterior cingulate cortex and medial prefrontal cortex), frontoparietal network (FPN, dorsolateral prefrontal cortex and inferior parietal lobule), and salience network (SN, anterior insula and dorsal anterior cingulate cortex) from 42 human participants using intracranial electroencephalogram (iEEG). We observed stronger within-network connectivity of the DMN, FPN and SN in broadband iEEG power, stronger phase synchronization within the DMN across theta and alpha bands, and weaker phase synchronization within the FPN in delta, theta and alpha band. We also found positive power correlations in high frequency band (70-170Hz) and negative power correlations in alpha and beta band for FPN-DMN and FPN-SN. Robust negative correlations in DMN-SN were found in alpha, beta and gamma band. These findings provide intracranial electrophysiological evidence in support of the network model for intrinsic organization of human brain and shed light on the way how the brain networks communicate at rest.

## Introduction

A network perspective has been emphasized to be crucial for a comprehensive understanding of the brain as an integrated system (Pessoa, 2014), and to concern integrative brain function (Sporns, 2010). From the first study of spatially organized networks in ongoing brain activity by using functional magnetic resonance imaging (fMRI; (Biswal, Yetkin, Haughton, & Hyde, 1995), in the past two decades, resting state networks (RSNs), defined as correlations of hemodynamic signals between brain regions at slow time scales in wakeful rest, have been extensively characterized by fMRI-based approaches (M. D. Fox & Raichle, 2007). Previous studies have identified at least three canonical networks: a) The default mode network (DMN) which mainly includes the posterior cingulate cortex (PCC) and medial prefrontal cortex (mPFC; (Raichle et al., 2001); b) The frontoparietal network (FPN) that including the dorsolateral prefrontal cortex (dlPFC) and inferior parietal lobule (IPL); c) The salience network (SN), including the anterior insula (AI) and dorsal anterior cingulate cortex (dACC) (Chen et al., 2013). These networks are thought to interact. For example, FPN and SN are consistently found to negatively regulate activity in the DMN (Fox et al., 2005), and are jointly involved in attention, working memory, decision making, and other high-level cognitive functions (Dosenbach et al., 2007; Aaron Kucyi, Jessica Schrouff, et al., 2018; Power et al., 2011). Noninvasive techniques such as fMRI have provided significant insights into the research of human brain networks, however, the electrophysiological basis in the brain network organization was still under-investigation.

Intracranial electroencephalography (iEEG) provides a means of electrophysiological investigations on RSNs in human participants. The extant iEEG studies have provided some initial electrophysiological findings for relationships among distinct brain regions that are typically thought as key nodes of brain networks, though the results seemed to be discrepant. For example, the PCC, one key region of DMN, has been reported to be correlated with the angular gyrus (Foster, Rangarajan, Shirer, & Parvizi, 2015) and mPFC (A. Kucyi et al., 2018) through high frequency band power fluctuations in resting state, while another study failed to find such high-frequency band DMN correlates but found stronger theta (4-8 Hz) band-limited power correspondence in the DMN and FPN, and stronger alpha (8-12 Hz) correspondence in the SMN and DAN (Hacker, Snyder, Pahwa, Corbetta, & Leuthardt, 2017). These discrepancies on the carrier frequencies of brain networks in electrophysiological signals may probably come from the small sample sizes and limited electrode coverage. However, it remained unclear, and no consensus for the role of frequencies has been emerged.

Here we reported a comprehensive iEEG investigation of activities and interactions among the DMN, FPN and SN in 42 participants with depth electrodes implanted directly within key nodes of these networks, aiming to probe the electrophysiological mechanisms of intrinsic organization of brain networks. It is now typically thought that there are two distinct types of intrinsic coupling modes. One type arises from phase coupling of band-limited oscillatory signals, the other results from coupled amplitude envelopes (or power) (Engel, Gerloff, Hilgetag, & Nolte, 2013). Therefore, in this study, to identify electrophysiological features of brain networks, we examined the correlates of networks from these two measurements, including both power correlations and phase synchronization. We hypothesized that 1) intra-network interactions would be significantly stronger than inter-network interactions, presenting as larger intra-network power correlations and phase synchronization; 2) network-to-network correlates showed frequency specific pattern, either in power correlations or phase couplings.

## Results

We obtained iEEG recordings of 3-minute rest from epilepsy patients in Shenzhen General Hospital. The limited electrode coverage of the iEEG recordings made it impossible to examine connectivity across the entire human brain with resolutions matching those of whole-brain fMRI studies. Therefore, we first selected electrode pairs that fell within the key regions of DMN (Default mode Network), FPN (Fronto-parietal Network), and SN (Salience Network) from individual brain, and then took an unbiased approach for assigning electrode pairs to brain networks which allowed us to probe network patterns in a common framework with fMRI studies. From a total of 7006 unique depth recording electrodes in 47 human participants, we selected 271 electrodes of 42 participants from regions void of pathological activity that fell within DMN/FPN/SN as defined in Yeo template (Yeo et al., 2011). Twelve participants had undergone fMRI scan during resting state, which helped us to identify the network pattern of selected regions with fMRI data. We first tested networks by calculating BLOD (blood oxygen level dependent, BOLD) correlations of 271 ROIs with the same locations in iEEG recording sites, and then assessed network connectivity patterns in iEEG signals with power correlations and phase synchronization (Figure 1).

**Figure 1.**
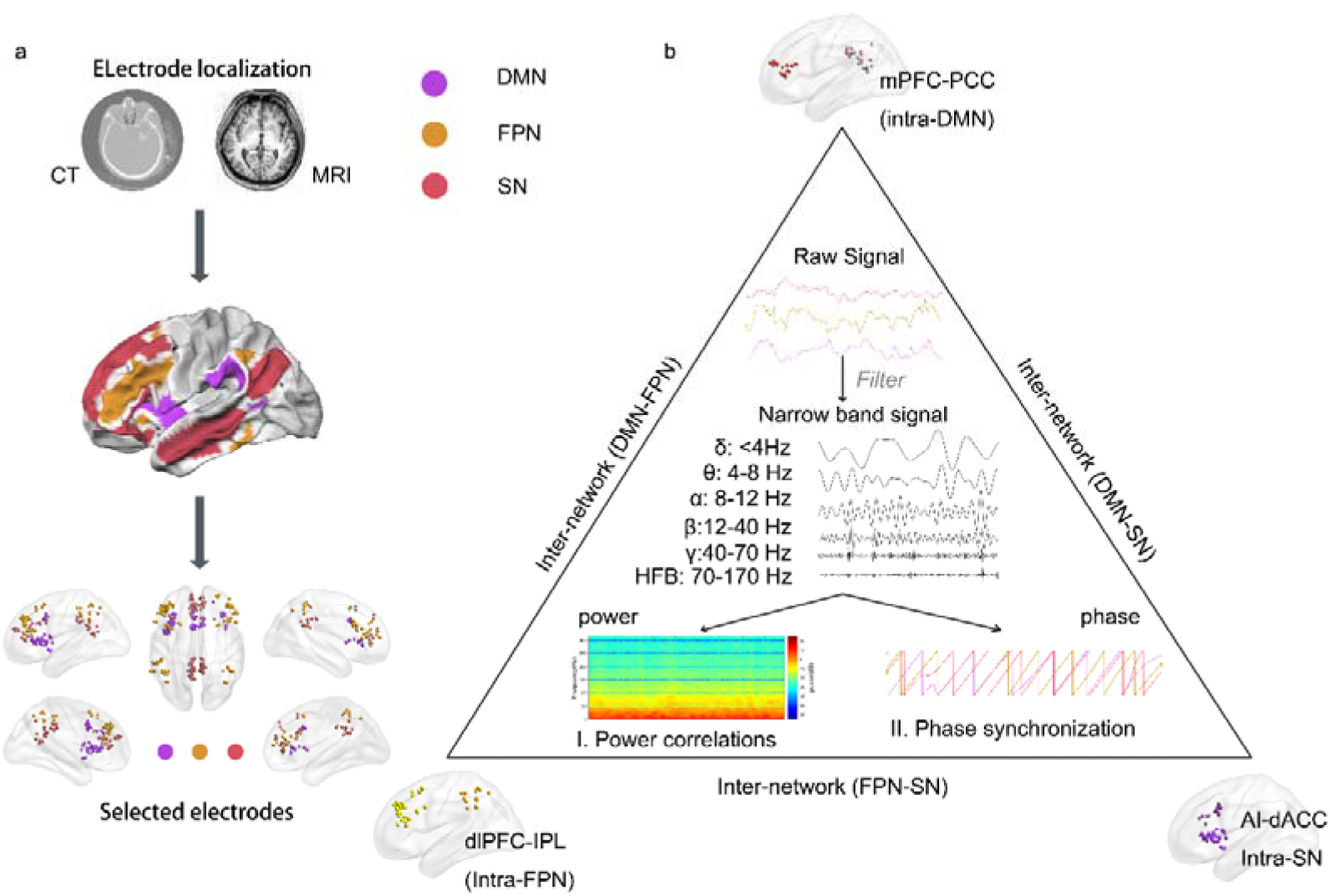
Schematic illustration of the methods. a) The top panel presents structural MRI images (T1) and CT scan images, co-registered to Montreal Neurological Institute (MNI) standard brain (MNI152_1mm_brain) to get MNI coordinate of each electrode. The middle panel presents Yeo template marked by 3 different colors denoted 3 networks (red for DMN, purple for SN and orange for FPN). The bottom panel shows all 271 selected electrodes that fell within DMN, FPN and SN. DMN, default mode network; FPN, frontoparietal network; SN, salience network. b) Schematic of connectivity estimation. Intra-network and inter-network connectivity were estimated by phase synchronization and power correlations.

### Stronger intra-network connectivity verified the existence of brain network with BOLD signals during resting state

Using each electrode location as a seed region, we calculated seed-based FC with all other electrode locations via Pearson correlation coefficient (*r*) of the resting state fMRI BOLD signals and converted to z using the Fisher-z transformation. To minimize the effect of distance on connectivity, based on the resting fMRI BOLD data, we created scatter plots of seed distances and BOLD *r-z* valuesWe constructed a curve by curve fitting the scatter plots and obtained a function that best describes their relations: BOLD *r-z* =1.295/distance (Figure 2a). We then removed the effect of distance on BOLD connectivity based on this fitting function.

**Figure 2.**
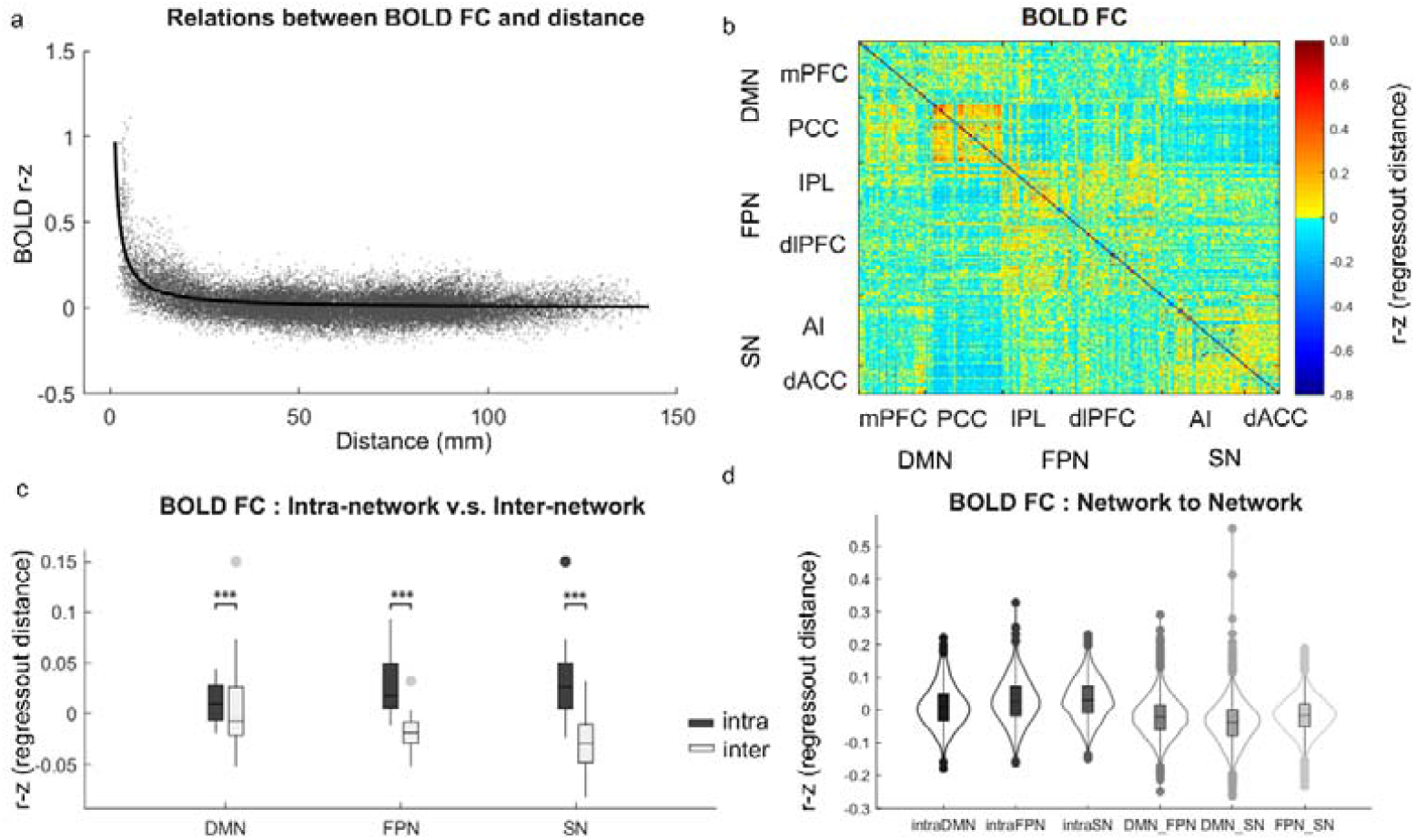
Resting state networks based on BOLD signals of selected 271 seeds. a) Relations between BOLD connections and distance of seed pairs. b) Correlation matrix of 6 regions of the DMN, FPN and SN based on functional connectivity of BOLD signals. BOLD correlations are averaged of 12 participants. c) Intra-network v.s. inter-network BOLD connections comparisons. Dark gray denotes intra-network, and light gray denotes inter-network. d) Between-network BOLD connectivity.

We averaged the BOLD correlations across 12 participants and obtained a correlation matrix (figure 2b) of 6 regions of these 3 networks (DMN/FPN/SN). The two-way ANOVA (3; DMN, FPN vs. SN) * (2; intra-network vs. inter-network) was applied to test the intra-/inter-connectivity differences among the DMN, FPN and SN. We found stronger intra-network connectivity than inter-network connectivity with a significant main effect on intra-/inter-connectivity (F=84.996, *p*<0.001, figure 2c). Note that no significant interaction (F=1.167, *p*=0.315), or main effect on network differences (F=2.441, *p*=0.092) were found, indicating that all the 3 networks (DMN/FPN/SN) could be recognized by fMRI BOLD signals, which further verified the validity of selected seed regions.

### iEEG intra- and inter-network connectivity

To estimate distance effect, we first calculated Pearson correlation coefficient *r* between distance and iEEG connectivity in power correlations and weighted Phase Lag Index (wPLI). We found that distance was significantly negatively correlated to power correlations (*r*=0.16, *p*<0.001; figure3a). Using linear regression with power coefficients *r* as dependent variable and distance as dependent variable, we obtained an equation that describes distance effect on iEEG power correlation coefficients: iEEG Power correlation coefficient (*r*)=-0.0014*distance+0.3605 We then removed distance effect based on that equation. We kept the original values of wPLI for phase synchronization as we did not find significant correlations between distance and wPLI (*r*=-0.053, *p*=0.168. Figure3b).

**Figure 3.**
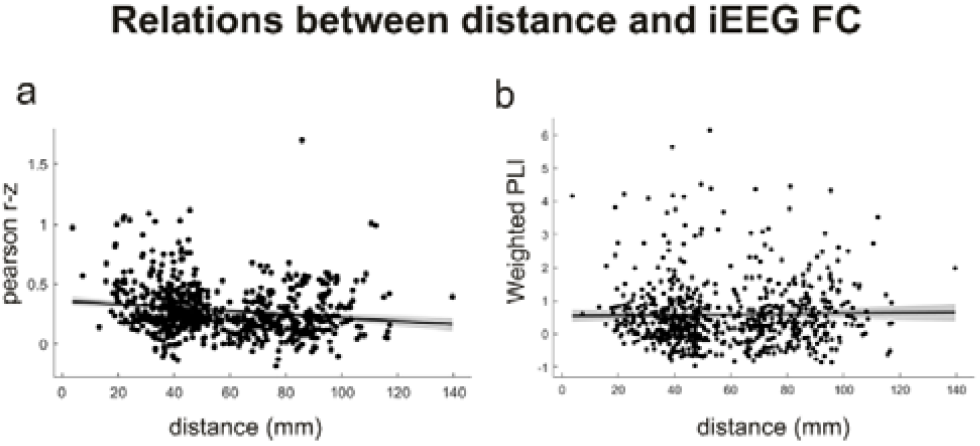
Relations between distance and iEEG FC. a) Relations between power correlations (measured as Pearson’s *r*, transformed to *z*) and distance. *r*=0.16, *p*<0.001. b) Relations between phase synchronization (measured as weighted PLI) and distance. *r*=-0.05 *p*=0.168.

Next, to assess whether there were specific features in intra-network patterns of each intracranial network, we conducted a 2 (intra-network / inter-network) * 6 (frequency bands) analysis of variance (ANOVA) on each network. We found that intra-network power correlations were significantly larger than that of inter-network (F[intra-DMN v.s. DMN-Others]=10.48, *p*=0.001; F[intra-FPN v.s. FPN-Others]=22.41,*p*<.001; F[intra-SN v.s. SN-Others]=7.83,p<.001; figure 4b), indicating that network could be recognized by broadband frequency power correlations of iEEG signals. There was no interaction effect with frequency (F[frequency*intra-DMN/inter-DMN]=1.77, *p*=0.116; F[frequency*intra-/inter-FPN]=0.11, p=0.332; F[frequency*intra-SN/inter-SN]=1.09, *p*=0.366). We also found specific greater across frequencies intra-network connectivity (power correlations) in each network of the DMN, FPN and SN (Figure 4b), identifying the intrinsic organization of intracranial networks.

**Figure 4:**
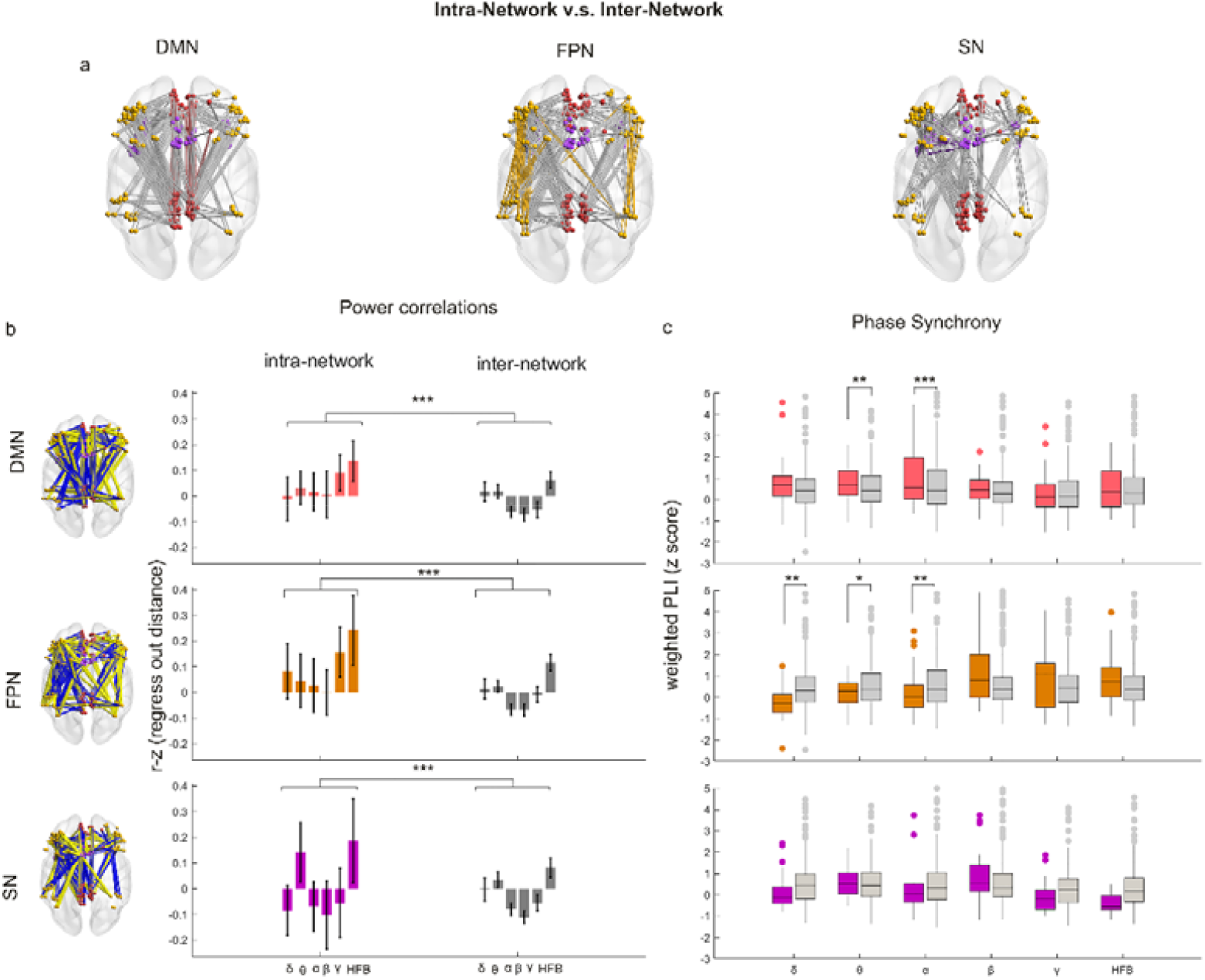
Intra-network v.s. inter-network connectivity based on iEEG signals. (a) Connections that included to the comparisons. Intra-DMN v.s. DMN-Others, intra-FPN v.s. FPN-Others, intra-SN v.s. SN-Others were presented from left to right, respectively. Colored lines denote intra-network connection, gray lines denote inter-network connection. (b) Intra-network v.s. inter-network connectivity comparisons based on power correlations. The left panel presents broadband power correlation patterns of DMN/FPN/SN (from top to down). Blue lines represent negative correlations, yellow lines represent positive correlations, thickness represents the degree of correlation. The right panel presents results of statistical test. X axis represents intra-network/inter-network, while y axis represents power correlation *r* (fisher-z transformed and regressed out distance). Top panel presents comparisons between intra-DMN and DMN-Others, middle panel presents intra-FPN and FPN-Others, and bottom panel presents intra-SN and SN-Others. delta, <4Hz; theta, 4-8Hz; alpha,8-12Hz; beta, 12-40Hz; gamma, 40-70Hz; HFB, 70-170Hz. ***, =*p*<.001). (c) Intra-network v.s. inter-network connectivity comparisons based on weighted Phase lag index (wPLI). X axis represents frequency bands, while y axis represents z values of PLI. Top panel presents comparisons between intra-DMN and DMN-Others, with red bars representing intra-DMN and gray bars representing DMN-others. Middle panel presents intra-FPN and FPN-Others, with orange bar representing intra-FPN and gray bar representing FPN-others. Bottom panel presents intra-SN and SN-Others with purple bars representing intra-SN and gray bars representing SN-other. *, *p*<.05; **, *p*<.01; ***, *p*<.001.

To determine whether there was network evidence for iEEG phase synchronization, similar 2 (intra-network / inter-network) * 6 (frequency bands) analysis of variance (ANOVA) were conducted to wPLI. We found that DMN and FPN can be recognized by iEEG frequency-specific phase synchronization (F[intra-DMN/DMN-Others*frequency bands]=2.62, *p*=0.023; F[intra-FPN/FPN-Others*frequency bands]=5.37, *p*<0.001), while SN not (F[intra-FPN/FPN-Others*frequency bands]=1.86, *p*=0.0985). Pair-wise comparisons showed that for DMN, PLI (θ/α intra-DMN) > PLI (θ/α DMN-Others), *p*(θ)=0.006, *p*(α)<.001; for FPN, PLI (δ/θ/α intra-FPN) < PLI (δ/θ/α FPN-Others), *p*(δ)=0.001, *p*(θ)=0.045, *p*(α)=0.004 (Figure 4c).Importantly, intra-DMN connectivity with wPLI in θ and α band were significantly stronger than connectivity of DMN-Others (Figure 4e).

### Frequency specific between-network iEEG power correlations

We then examined the oscillational mechanisms between brain networks. One-sample *t* test was used for inter-network (DMN-FPN, DMN-SN, FPN-SN) power correlations. We found that DMN and FPN were negatively correlated in α and β band, but positively correlated in HFB (*p* (alpha)=0.001, *p*(β)=0.028, *p*(HFB)<0.001, FDR corrected; figure 5), indicating that DMN and FPN were anti-correlated in alpha and beta band and correlated in HFB. We observed negative correlations in alpha, beta, and gamma band between DMN and SN (all *p*s<0.001, FDR corrected), indicating that DMN and SN were anti-correlated (figure 5). FPN and SN were negatively correlated in α and β band, and positively correlated in θ band and HFB (One-sample *t* test, *p*(α/β/HFB)<0.001, *p*(θ)=0.045). Overall, these results reflected that inter-network connectivity based on iEEG power correlations was frequency dependent.

**Figure 5:**
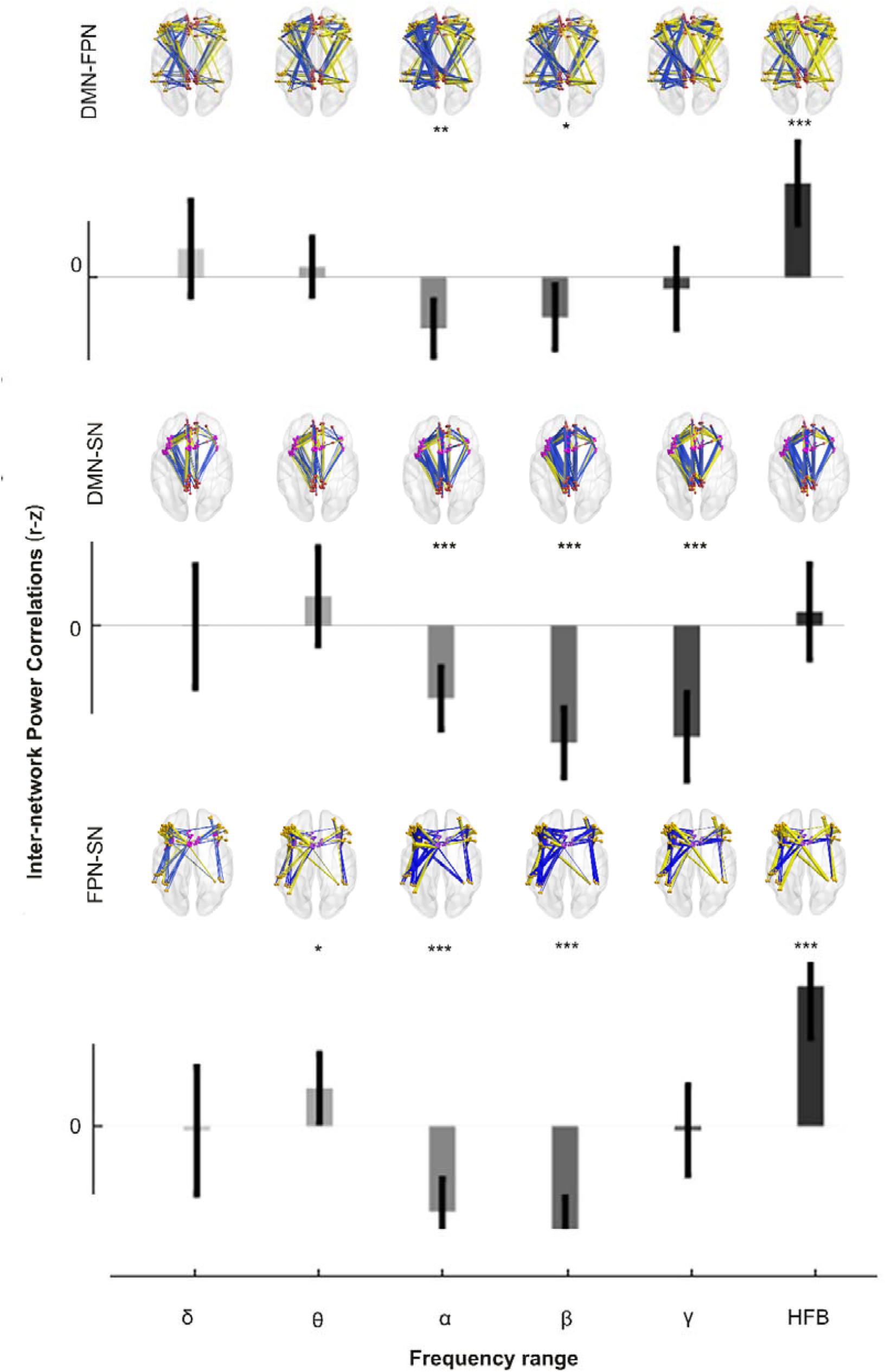
Inter-network connectivity based on iEEG power correlations. Inter-network connections in each frequency band (delta, theta, alpha, beta, gamma, and HFB) based on iEEG power correlations. The upper panel is connectivity between DMN and FPN, the middle is between DMN and SN, and the bottom panel is between FPN and SN. Blue lines represent negative values and yellow represent positive ones. Thickness represents correlation degree. Note that only the significantly connected electrodes were shown. X axis represents frequency bands, y axis represents *z* values of power correlation *r*. One sample *t* test was applied to test the significance to connectivity. *, =*p*<.05; **, *p*<.01; ***, *p*<.001; FDR corrected.

## Discussion

In this study, we examined oscillational mechanism of intrinsic brain networks based on intracranial recordings. For technical reasons and electrodes coverage limitation, we did not analyze the data at individual level but unbiasedly assigned electrode pairs to brain networks We evaluated intrinsic coupling modes between and within networks using power correlations and phase synchronization. We found that broadband frequency power correlations of iEEG signals distinguished DMN, FPN and SN, while frequency-specific phase synchronization separated DMN and FPN. Our findings provide novel evidence for frequency-dependent segregation and integration on the organization of the brain networks at rest.

### 1. Broadband frequency power correlations separate brain RSNs

In line with prior work on RSNs of BOLD signals in human brain (Yeo et al., 2011) and electrophysiological characteristics of RSNs in macaque monkeys (Liu, Yanagawa, Leopold, Fujii, & Duyn, 2015), our results show distinctively intracranial networks (DMN, FPN and SN) with broadband power correlations in human. Together with previously reported positive correlations of broadband power with BOLD amplitude (Hermes, Nguyen, & Winawer, 2017), these results based on BOLD signals of fMRI studies jointly reveal electrophysiological mechanisms of the RSNs. Specifically, for example, theta (4-8Hz) band power is associated with fluctuations of the DMN and FPN, alpha (8-12Hz) relates to activity of the SMN and DAN (Hacker et al., 2017), and within-network coupling has been observed in the HFB range (A. Kucyi et al., 2018). However, a recent iEEG study did not show any correspondence in the DMN by comparing power spectral density in different frequency ranges and high gamma broadband power correlation between intra- and inter-network connectivity (A. Das, de Los Angeles, & Menon, 2022). Because they did not compare intra-/inter-power correlations in the broadband signals but in the band-limited range, it not necessarily contradicted to our results that broadband power correlations separated distinct networks.

### 2. Frequency dependent between-network connectivity

Focusing on network interactions, we show that iEEG power correlations of the DMN was negatively correlated with those of the FPN and SN in α/β band while positively correlated with those of the FPN in high gamma band, indicating a frequency-dependent between-network relationship. The results support the proposed mechanism of functional brain that populations of neurons transmit information by coordinating their oscillatory activity with the receptor oscillations at certain frequencies and different frequencies subserve different functions (Buzsáki & Draguhn, 2004). Previous fMRI studies have shown that the DMN exhibits infra-slow anticorrelated activity with the FPN and SN across various behavioral states including wakeful rest (Michael D Fox et al., 2005; Fransson, 2006; Aaron Kucyi, Amy Daitch, et al., 2018), however, the antagonistic relationship of the DMN with FPN and SN has been contentious because distributions of resting state BOLD correlations are heavily skewed towards positive values (Keller et al., 2013; Murphy & Fox, 2017). Our findings suggest that the DMN is not always negatively correlated with FPN or SN at rest, which is dependent on communicating frequencies. The controversial antagonistic relationship from fMRI findings might due to the BOLD signal is an indirect measure related to hemodynamics, which could not comprehensively reflect the electrophysiological interactions between spatially separated networks.

Unfortunately, specific cognitive function of different oscillatory frequencies was not examined in the current study. Connectivity in various frequency bands have been shown to paly different functional roles (Hahn et al., 2019). For example, alpha-band oscillations have been associated with functional inhibition (Jensen & Mazaheri, 2010); oscillations or couplings in the beta-band are related to maintenance of the current cognitive state (Engel & Fries, 2010). Taken together, negative correlations between the DMN and FPN/SN in alpha/beta band in our study may speculatively suggest that the DMN and FPN/SN are two competing systems in maintaining a cognitive state. To this end, future studies with specific task manipulation is necessary.

### 3. Frequency specific phase synchronization

Phase relations, in the form of phase-modulated neuronal firing, has largely been identified in rodents, monkeys and humans (Maris, Fries, & van Ede, 2016). Phase relation diversity has been thought to support information transmission between two brain regions and allow for concurrent segregation of multiple information streams (Fries, 2005). Our findings suggest a frequency specific phase correlated pattern for the DMN and FPN. Specifically, stronger theta (4-8Hz) and alpha (8-12Hz) phase synchronization in the DMN and weaker phase synchronization in low frequency range (delta, theta and alpha) in the FPN, suggest phase coding as a property for functional communications in specific network. Long-range phase synchrony in the α-oscillation band has been proposed to facilitate information integration across anatomically segregated regions (Sadaghiani et al., 2012), in support of attentional, executive, and contextual functions (Palva & Palva, 2011). Therefore, these results may suggest the active role of the DMN rather than the FPN to maintain our cognitive functions at rest. Theta phase synchronization, especially in the hippocampus of the DMN, has been largely found in memory system in both animals (Wikgren, Nokia, & Penttonen, 2010) and humans (Clouter, Shapiro, & Hanslmayr, 2017), while the mPFC of the DMN has been shown to be phase coupled during memory processing (Kaplan et al., 2014). Our results suggest the active theta coupling of the DMN at rest, which may contribute to maintenance of the memory function. Interestingly, a recent study observed greater intra-DMN phase synchronization in the slow-wave (<4Hz) while greater cross-network interaction of the DMN in higher frequencies (>4Hz; Anup Das, de los Angeles, & Menon, 2020). Electrodes in temporal regions, such as superior temporal sulcus (STS) and middle temporal gyrus (MTG) consisted most of the DMN electrodes included in their study, which is highly different from our study as we focus more in medial prefrontal cortex (mPFC) and posterior cingulate cortex (PCC). This might be a potential reason for the inconsistency.

## Limitations

One most serious limitation for this study is the nature of the patient population. All the data were collected from intractable epilepsy patients. Therefore, whether the current findings and conclusions can be generalized to other populations is unknown, even though we avoid recordings from the region that had been related to the epilepsy by the presence of interictal discharges, structural lesions or evidence of involvement in seizure onset. Another limitation of the current study might be the lack of simultaneous behavioral measurements of the cognitive functions while iEEG being recorded, which would be necessary in the future study.

To conclude, we identify the classical resting state networks of DMN, FPN and SN by using intracranial electrophysiological recordings, supportive of the network perspective on the intrinsic organization of our brain. We show the frequency-specific way of the network communications at rest. Our results highlight the importance of identifying the carrier frequency that underlies communications within and between networks. Our work shed light on the understanding of neurophysiology of the brain network organization.

## Methods

### Participant selection

There were 47 patients with drug-resistant focal epilepsy, undergoing presurgical assessment with depth electrodes stereotactically for clinical evaluation participated the study at Shenzhen General Hospital. We selected data of the participants based on the inclusion criteria that each participant should have electrodes located at least two nodes of the six regions of interest (ROIs), including the posterior cingulate cortex (PCC), medial prefrontal cortex (mPFC) within the default mode network (DMN); the dorsal lateral prefrontal cortex (dlPFC) and inferior parietal lobule (IPL) within the frontoparietal network (FPN); the dorsal anterior cingulate cortex (dACC) and anterior insula (AI) within the salience network (SN).

Finally, data from 42 participants were included in the data analyses (age: 29.93 ± 8.00 (*Mean* ± *SD*), 16 females and 26 males, all right-handed. See sup.tabel for details in demographic information. Twelve participants (5 females and 7 males, age: 29 ± 8.60 (*Mean* ± *SD*)) had undergone resting-state fMRI scans which was used to assess the validity of selected regions in forming brain networks.

### SEEG data acquisition

After electrodes implantation, patients were monitored over several days while awake and comfortable in their hospital rooms. During this period, we collected their SEEG data at rest.

We acquired local field potentials (LFPs) from brain tissues with platinum-iridium, multi-lead electrodes in either 256 or 192 channel EEG amplifier system (NIHON-KOHDEN NEUROFAX EEG-1200) with a sampling rate of either 2000 or 2500 Hz. Each electrode was 2mm long, 0.8 mm thick, and inter-electrode spacing was 3.5mm. The neuroanatomical targets and numbers of electrodes implanted to each participant varied exclusively according to clinical requirements (for details, see sup.table 1). Before each 3-minute set of resting-state recording, participants were instructed to stare at a fixation cross with eyes open and do not think of anything specific.

### MRI acquisition

Pre-operative MRI scanning were conducted on a 3T SEIMENS Trio scanner equipped with a 32-channel head coil. During resting state fMRI, participants were instructed to close their eyes but remain awake during the eyes closed condition. The fMRI scan using the 32-channel phased-array head coil supplied by the vendor. High-resolution three-dimensional T1-weighted magnetization prepared rapid acquisition gradient echo images were acquired for anatomic reference [repetition time (TR) =1900 ms; echo time (TE) = 2.2 ms; flip angle (FA) = 90°; 1.0 mm isotropic voxels]. Resting-state fMRI were acquired through EPI (echo planar imaging, EPI) sequence [TR = 2000 ms; TE = 30 ms; FA = 90°, 3.6 mm isotropic voxels; 240 volumes in 8 minutes]. In addition, a computed tomography (CT) scan was obtained following electrode implantation, which was used for anatomical localization of electrode contacts.

### MRI preprocessing

The preprocessing of functional image were performed using Data Processing and Analysis of Brain Imaging (DPABI) (Yan, Wang, Zuo, & Zang, 2016). Steps were as follows: First, temporal and spatial corrections were performed, including slice time and head motion correction, furthermore normalized (voxel size: 3□mm) into EPI template. Any participants who had a maximum translation in any of the cardinal directions larger than 3□mm or a maximum rotation larger than 3° were excluded from subsequence analysis. Second, detrending analysis was performed on the normalized data to minimize the effect of linear trend. Third, nuisance signals were regressed out from functional image through linear regression analysis. The nuisance signals included six motion parameters and their first temporal derivative, white matter and cerebrospinal fluid signals. Additionally, the global signal (GS) averaged over the whole brain and its temporal derivative were removed by linear regression.

### SEEG preprocessing

SEEG recordings was preprocessed with publicly available tool Fieldtrip (Oostenveld, Fries, Maris, Schoffelen, & neuroscience, 2011). Signals were first down-sampled to 1000Hz. Then all channels were visually inspected; noisy and flat channels exceeding 5 standard deviations above or below the mean across channels were rejected, as well as channels with abnormal or interictal spikes. Notch filter was performed at a bandwidth of 4Hz with ‘butterworth’ approach to remove line noise at 50Hz and its harmonics. The iEEG data were re-referenced to a common average. To reduce boundary and carry over effects, we discarded 500 sample points of each iEEG data from the beginning.

### Electrode localization

Positions of depth electrodes were localized by pre-operative structural magnetic resonance imaging (MRI) scans and post-operative computed tomography (CT) scans using an in-house developed software. Structural MRI images were recorded before implantation, and rigid-body co-registration was used to colocalize MRIs and postimplant CT scans. Individual MRI images were registered to a Montreal Neurological Institute (MNI) standard brain (MNI152_1mm_brain). The coordinates were converted to standard MNI space using the transformation matrix. Based on the MNI coordinates, we selected 271 electrodes of interest that fell within the DMN, FPN and SN, as defined in Yeo cortical atlas (Yeo et al., 2011). Depth electrodes included in the following analysis were shown in figure 1b.

### fMRI functional connectivity

Using the electrode coordinates in fMRI volume space, we extracted the time-series from a 2 mm radius sphere at each electrode location. We adopted a classical “seed-based” functional connectivity (FC) analysis, where the time course recorded at a selected seed region was correlated with those at other target locations. We defined the electrode contacts as seed region and calculated its BOLD correlations with all other electrode contacts using Pearson correlation coefficient (*r*) of time series. Each *r* value would be converted to *z* value by fisher transformation for the subsequent analysis.

### iEEG Power correlations

To obtain reliable spectral estimates, we utilized a proach (Thomson, 1982) based on discrete prolate spheroidal sequences (DPSS). Codes were adapted from a previous study (Prerau, Brown, Bianchi, Ellenbogen, & Purdon, 2017). Parameters were set as 9 tapers for 5s segments, step in 1s, frequency smoothing of ±1*Hz* ranging from 0-170Hz. The obtained power spectrum within the following 6 frequency bins (delta: <4Hz; theta: 4-8Hz; alpha:8-12Hz; beta: 12-40Hz; gamma: 40-70Hz; HFB: 70-170Hz) were averaged for the subsequent analyses. Power correlations were estimated via Pearson’s correlation coefficient (*r*) of power time series between electrode pairs. To satisfy test assumptions, we normalized *r* values by Fisher’s z transformation.

### iEEG Phase synchronization

We calculated weighted Phase Lag Index (wPLI) as the index of phase synchronization between two time-series proposed by Vinck et al.(2011), which shows that WPLI outperforms PLI, coherence, and imaginary coherence (IC) with real local field potentials (LFP) data. For given time-series X(*t*) and Y(*t*) at time point t, PLI is defined as the absolute value of the sum of the signs of the imaginary part of the complex cross-spectral density *S*_*xy*_, while wPLI weights the cross-spectrum according to the magnitude of the imaginary component, which allows it to limit to the influence of cross-spectrum elements around the real axes which are at risk of changing their “true” sign with small noise perturbations:

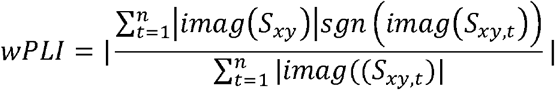

Here, wPLI was computed using scripts adapted by Fieldtrip toolbox based on MATALB (version 2017b). The mean value across the frequency-bins (steps at 1Hz) in the frequency range (delta: <4Hz; theta: 4-8Hz; alpha:8-12Hz; beta: 12-40Hz; gamma: 40-70Hz; HFB: 70-170Hz) of interest was calculated to obtain a single wPLI coupling value. We constructed surrogate wPLI values by shuffling labels. This procedure was repeated 1000 times to build a distribution of surrogate values of wPLIs at each frequency across electrode pairs. A z-score was obtained by subtracting the average of the surrogate data and dividing the result by the standard deviation of the surrogate distribution. We averaged the surrogate PLI values across channels in order to get the estimates of the surrogate means and standard deviations.

### Intra- and inter-network comparison

Both iEEG and BOLD correlations are systematically greater at short distances, thus prior to compare iEEG intra-network and inter-network connectivity, we performed additional control analysis to rule distance out as a potential factor for the higher intra-network connectivity. We calculated the Euclidean distance between all included electrode pairs and estimated the effectiveness of distance on connections by curve fitting equation.

Intra-network connectivity was assessed between all electrode pairs across distinct brain regions within the selected network (DMN/FPN/SN). To minimize potential bias from sampling of electrodes within the same brain region, we did not include same-region electrode pairs. For example, for intra-DMN connectivity analysis, we included electrode pairs of PCC-mPFC but not PCC-PCC. Inter-network connectivity was assessed between all electrode pairs across networks. For example, for inter-DMN electrode pairs, one electrode located in the DMN, and the other either in the FPN or SN. We would be able to identify distinctive networks if intra-network connectivity had specific features different from that of inter-network.

## Data sharing

The datasets reported in the current study and data analysis scripts are available upon request from corresponding author.

## Acknowledgement

This work was supported by the National Natural Science Foundation of China (31871137, 31920103009 and 32071100), the Major Project of National Social Science Foundation (20&ZD153), Young Elite Scientists Sponsorship Program by China Association for Science and Technology (YESS20180158), Guangdong International Scientific Collaboration Project (2019A050510048), Natural Science Foundation of Guangdong Province (2020A1515011394), Shenzhen-Hong Kong Institute of Brain Science-Shenzhen Fundamental Research Institutions (2019SHIBS0003), Shenzhen Science and Technology Research Funding Program (JCYJ20180507183500566 and JCYJ20190808121415365).

